# Phage-Encoded Bismuth Bicycles: Instant Access to Targeted Bioactive Peptides

**DOI:** 10.1101/2024.04.10.588800

**Authors:** Sven Ullrich, Upamali Somathilake, Minghao Shang, Christoph Nitsche

## Abstract

Genetically encoded libraries play a crucial role in discovering structurally rigid, high-affinity macrocyclic peptide ligands for therapeutic applications. This study represents the first genetic encoding of peptide-bismuth and peptide-arsenic bicyclic peptides in phage display. We introduce bismuth tripotassium dicitrate (gastrodenol) as a water-soluble Bi(III) reagent for phage library modification and *in situ* bicyclic peptide preparation, eliminating the need for organic co-solvents. Additionally, we explore As(III) as an alternative thiophilic element used analogously to our previously introduced class of peptide-bismuth bicycles. The modification of phage libraries and peptides with these elements is instantaneous and entirely biocompatible, offering an advantage over conventional alkylation-based methods. In a pilot display screening campaign aimed at identifying ligands for the biotin-binding protein streptavidin, we demonstrate the enrichment of bicyclic peptides with dissociation constants two orders of magnitude lower than those of their linear counterparts, underscoring the impact of structural constraint on binding affinity.

## Introduction

Genetically encoded libraries have risen to key technologies in peptide drug discovery.^1-3^ Integrated into display screening platforms, their extensive sequence diversity facilitates the identification of peptide hits for therapeutically relevant targets.^4-6^ Chemical library modifications compatible with the biological environment of these screening platforms have been shown to enhance the pharmaceutical properties of the displayed peptides.^7-9^ Particularly desirable are strategies that constrain the entire peptide structure, such as macrocyclisation or bicyclisation.^9-11^ Restricting the conformational freedom of a peptide can cause dramatic improvement of bioactivity, stability and membrane permeability.^9,12,13^ Hence, constrained peptides derived from genetically encoded library selections are anticipated to initiate a new wave of peptide therapeutics.^3,14-16^

Peptide-bismuth bicycles are an emerging class of constrained peptides.^17,18^ They feature a unique bicyclic architecture, with the peptide component connected to a trivalent bismuth core *via* three sulphur atoms from cysteine residues.^17^ We have recently demonstrated that the bicyclisation of peptides through bismuth leads to improved bioactivity, metabolic stability and cell permeability compared to their linear counterparts.^17,18^ Importantly, the modification occurs instantaneously and can be initiated *in situ*.^17^ Furthermore, the radioisotope bismuth-213 is an important emitter in targeted alpha therapy,^19,20^ highlighting the potential for peptide-bismuth bicycles in future cancer therapy. With increasing demand for bicyclic peptides as an innovative therapeutic modality,^11,14,16,21-23^ we set out to investigate the compatibility of bismuth bicyclisation with genetically encoded libraries, specifically phage display, to allow for the rapid identification of peptide-bismuth bicycles for virtually any given target.

Phage display is a versatile screening technique with extensive application in antibody development and drug discovery.^24-26^ It requires the presentation of peptides on the M13 bacteriophage surface, e.g., on the coat proteins pIII or pVIII (**Figure 1a**), which is facilitated by the integration of a semi-randomised library into the phage genome.^24-26^ Chemical modification of the displayed peptides expands library diversity beyond the standard genetic code, enabling the selection of peptide ligands with enhanced affinity and stability, including macrocyclic peptides.^7,14,16,27^ Conventional approaches for the display of bicyclic peptide libraries on phage centre on the modification of three cysteine residues using organic alkylating agents.^7,28^ Most commonly, 1,3,5-tris(bromomethyl)benzene (TBMB) is used (**Figure 1b**),^28^ despite a multitude of inherent disadvantages of the reagent concerning genetically encoded systems. Most notable is the compromised selectivity of TBMB, as non-thiol nucleophiles as well as additional cysteine residues in peptides and proteins may cross-react.^28,29^ Likewise, the reagent is incompatible with the commonly used thiol-free reducing agent tris(2-carboxyethyl)phosphine (TCEP).^28^ Furthermore, TBMB typically requires higher temperatures and prolonged incubation periods for sufficient modification.^28^ Additionally, it mandates extensively diluted phage solutions and significant amount of organic co-solvent due to its limited buffer solubility (**Figure 1b**).^28^ Nevertheless, TBMB and its derivates have contributed considerably to the successful identification of bicyclic peptide ligands for various protein targets.^14^

**Figure 1.**
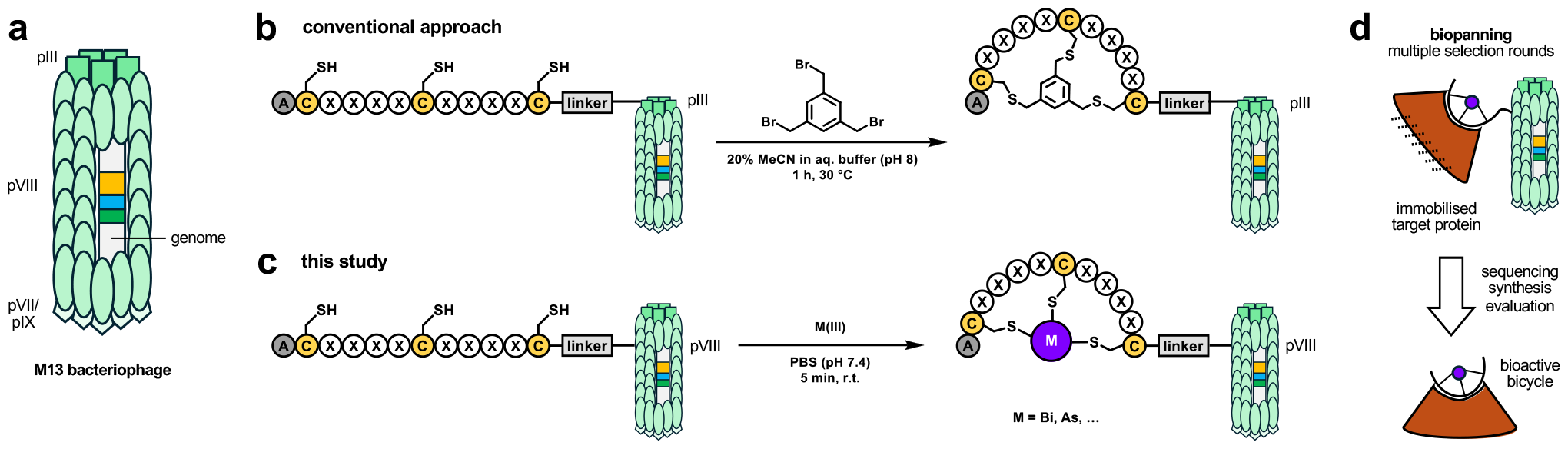
(a) Schematic representation of the M13 bacteriophage structure used in this study. (b) Conventional approach to display bicyclic peptide libraries on pIII of the M13 phage. (c) Our newly developed strategy to display bicyclic peptide libraries on pVIII of the M13 phage using thiophilic metals like bismuth(III) or arsenic(III). (d) Panning and selection of modified phage libraries against protein targets to identify high-affinity bicyclic peptides with a metal core.

In contrast to TBMB, the reaction of three cysteines residues in a peptide with bismuth(III) is instantaneous, selective and proceeds in aqueous buffer at room temperature and physiological pH.^17^ We therefore hypothesised that the strong selectivity and biocompatibility of bismuth-based peptide bicyclisation will be a valuable technique in the context of phage display (**Figure 1d**), where it could also mitigate the limitations of commonly used alkylating agents. In addition, we demonstrate for the first time that the concept of peptide bicycle formation by bismuth can be extended to other thiophilic metals of interest (**Figure 1c**).

## Results

We chose the commonly used M13 phage for our investigations and selected the pVIII coat protein for bicyclic peptide display (**Figure 1a**). A semi-randomised library containing an ACX_4_CX_4_CGGGENLYFQS extension (where X is any canonical amino acid) at the N-terminus of pVIII formed the basis of our studies. This construct comprises a glycine linker and TEV protease recognition site between the displayed peptide and pVIII to enable selective peptide release from the phage if required. The equal spacing of cysteine residues in our construct is reminiscent of the libraries used for modification with TBMB and our previously published peptide-bismuth bicycles.^17,18,28^

Our initial considerations related to the modification of the phage library in PBS (pH 7.4) under fully biocompatible conditions. We employed bismuth tribromide (BiBr_3_) to replicate the conditions used for bismuth bicyclisation of peptides in our previous studies. Given that BiBr_3_ and most other Bi(III) salts are insoluble in water at near-neutral pH, concentrated DMSO stocks of bismuth salt are necessary for peptide modification in buffer solutions. Anticipating potential challenges associated with this method when applied to phages, we explored the fully buffer-soluble bismuth tripotassium dicitrate (gastrodenol) as an alternative. Consequently, we conducted screenings with both bismuth reagents to assess their efficacy in phage display. Thirdly, we envisioned As(III) from water soluble sodium arsenite (NaAsO_2_) as a substitute for Bi(III) in library modification, and used it analogously for phage library modification.

We used the biotin-binding bacterial protein streptavidin as a model target for screening. Our bicyclic peptide phage libraries were employed for selection against streptavidin immobilised on magnetic beads, involving four rounds of biopanning. We decided to conduct competitive elution with biotin in each round of the selection process to strongly bias the selection for peptides that bind to the ligand-binding site (**Figure 2a**). In addition to both Bi(III) screenings (BiBr_3_ and gastrodenol), we performed two As(III) screenings with NaAsO_2_ in order to assess the robustness and reproducibility of our strategy. Gratifyingly, both Bi(III) screening campaigns featured similarly enriched peptides despite the differences in the modification reagent (**Figure 2b–c**). The two separately conducted As(III) screenings yielded hits distinct from the Bi(III) results; however, enriched identical sequences, signifying excellent replicability (**Figure 2d–e**). The top hits contained an HPQ/M motif, which has previously been described to facilitate peptide interactions with the biotin binding site of streptavidin.^30-33^

**Figure 2.**
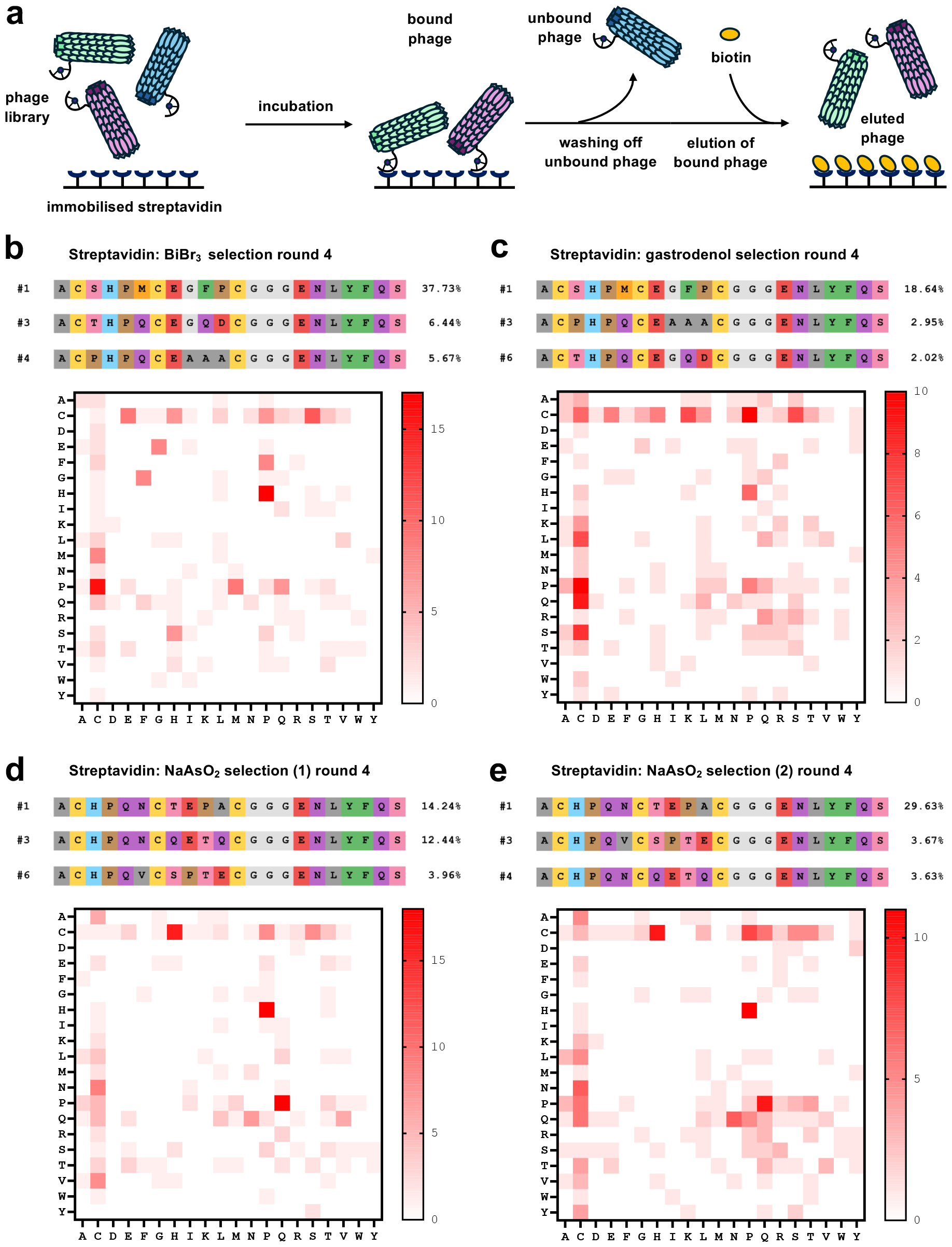
(a) Biopanning of a bicyclic peptide phage library against immobilised streptavidin using competitive elution with biotin. (b–e) Selected top hits identified by Nanopore sequencing of selection round 4 including dipeptide motif analysis of the flanked randomised part (CX_4_CX_4_C) of the top 25 peptide insert hits, which are provided in **Figures S6–S9**.

We synthesised four linear peptide sequences (H-ACX_4_CX_4_C-NH_2_) discovered in the Bi(III) and As(III) selection campaigns against streptavidin using automated solid-phase peptide synthesis (SPPS) on Rink amide resin. The purified peptides were bicyclised with water-soluble Bi(III) from gastrodenol and As(III) from NaAsO_2_ (**Figure 3a**). Bicyclisation is instantaneous and quantitative in buffer at physiological pH (**Figure 3b**). This *in situ* modification facilitated the direct evaluation of bicyclic peptides by SPR against immobilised streptavidin, eliminating the need for laborious purification of the bicyclic peptide. All four tested bicyclic peptides exhibited micromolar affinity towards streptavidin (**Figure 4**), with peptide **2b** from the Bi(III) screening campaign standing out as the best of the four analysed peptides (K_D_ = 7.4 µM). We selected the two peptides with the highest affinity from the bismuth (**2b**) and arsenic (**3a**) series and compared them to their linear analogues (**Figure 4**). Importantly, in both cases, the linear unmodified peptide exhibited significantly lower affinity for streptavidin. Arsenic bicycle **3a** is approximately 80 times more active than its linear counterpart **3**, while bismuth bicycle **2b** is even approximately 200 times more active than its linear precursor **2**.

**Figure 3.**
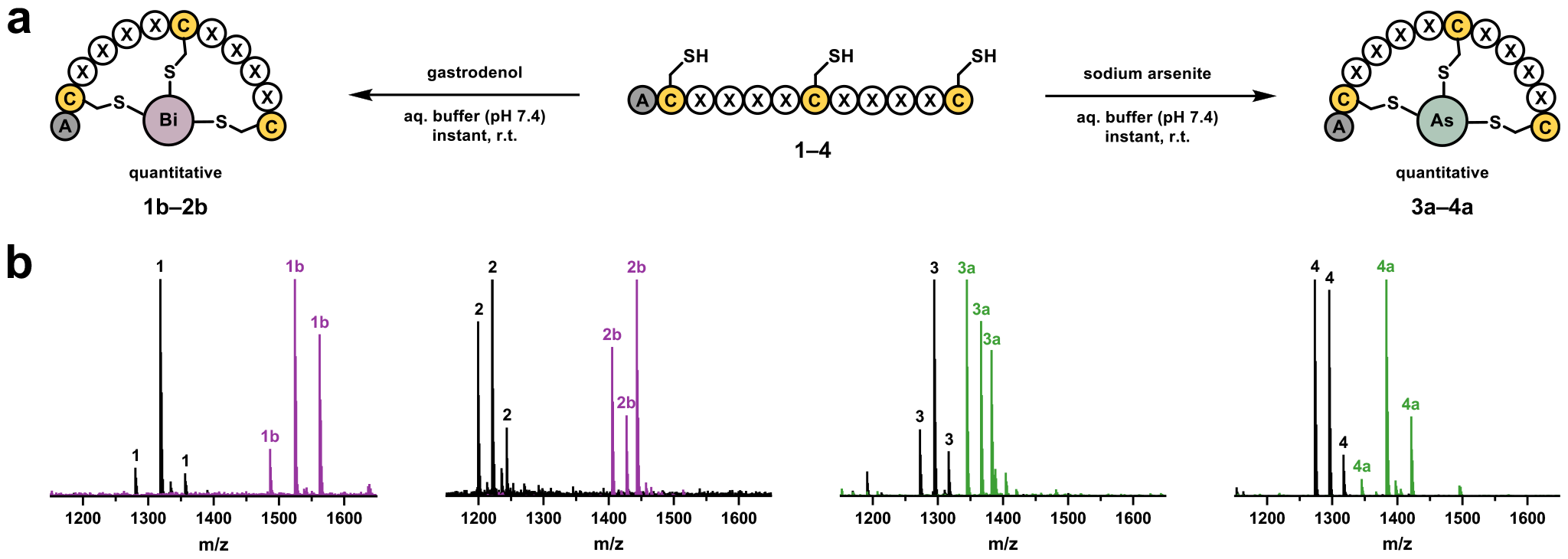
(a) Conversion of synthetic linear peptides from the streptavidin screening campaigns into peptide-bismuth bicycles (left) using gastrodenol (bismuth tripotassium dicitrate) or peptide-arsenic bicycles (right) using sodium arsenite (NaAsO_2_). (b) High-resolution mass spectrometry overlays of linear peptides (**1–4**) and their quantitative conversion into the respective bicycles (**1b– 2b, 3a–4a**). Specific mass adducts for each peak are indicated in **Figures S10 & S11**.

**Figure 4.**
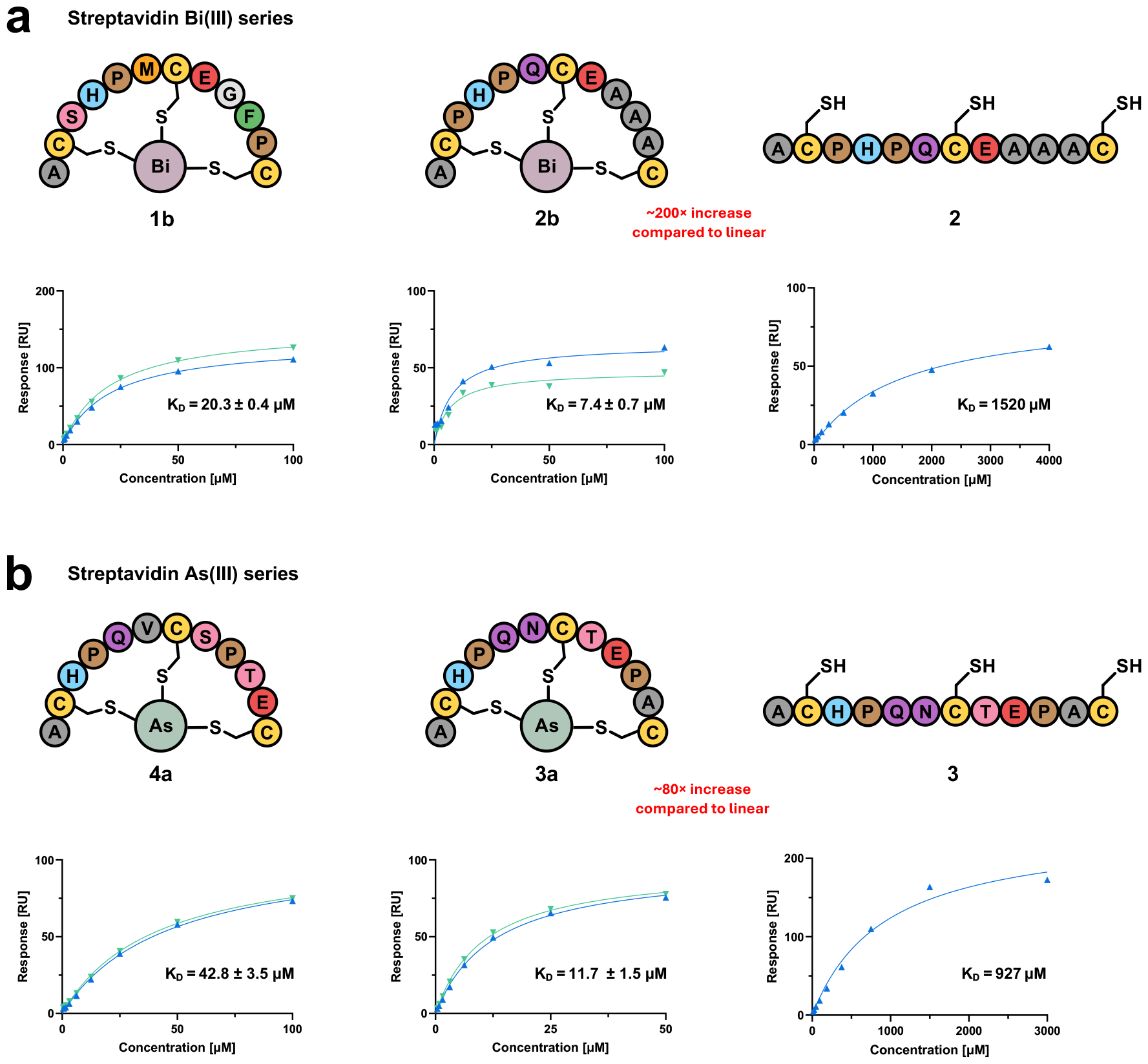
SPR evaluation of the interaction between selected bicyclic and linear peptides and streptavidin from the (a) bismuth (**1b, 2b, 2**) and (b) arsenic (**4a, 3a, 3**) screening series. Bicyclic peptides were tested in duplicate (green, blue), with their respective K_D_ values shown ± SD.

## Discussion

As recently demonstrated, linear peptides containing three cysteine residues can be modified by Bi(III) to generate constrained bicyclic peptides under biocompatible conditions.^17,18^ While our previous studies relied on BiBr_3_ dissolved in DMSO, in this study, we assess the pharmaceutical drug gastrodenol (bismuth tripotassium dicitrate) as an alternative easily accessible and water-soluble agent for the quantitative generation of bismuth-peptide bicycles from linear precursors in buffer at physiological pH. Additionally, we induced bicycle formation by As(III) using water-soluble NaAsO_2_ (**Figure 3**). While the formation of arsenic-peptide bicycles might be of limited interest for applications in biomedical research due to their expected toxicity, this example demonstrates the potential of expanding our macrocyclisation strategy to other thiophilic metals for theranostics. Modification of M13 phage libraries by gastrodenol and NaAsO_2_ offers notable advantages over conventional modification with TBMB because these reagents (i) eliminate the use of organic co-solvent, (ii) react instantaneously at room temperature, (iii) tolerate the reducing agent TCEP, and (iv) exhibit strong selectivity for the three cysteine residues.

We validated our methodology with screening campaigns against streptavidin as a model system. The remarkably strong interaction between streptavidin and biotin is extensively studied,^34^ and peptides binding to the biotin-binding site have been previously identified using phage display and alternative technologies.^30-33,35-39^ As a result, it is well-known that the motif HPQ/M is highly conserved in peptides binding to the biotin-binding site.^32,33,40^ Crystal structures with HPQ peptides are even documented (e.g., PDB: 1SLG, 1SLD).^32,41,42^ Competitive elution of phage with biotin ensured that most eluted phages bound to the biotin binding site. Consequently, all four of our screening campaigns displayed strong enrichment of the HPQ/M motif, as demonstrated by the dipeptide motif analyses conducted for each library (**Figure 2b–e**). Interestingly, both Bi(III) libraries exhibited the highest enrichment for an HPM peptide, while both As(III) libraries showed highest enrichment for an HPQ peptide. The fact that we obtained identical top hits from both arsenic and bismuth campaigns underscores the robustness and effectiveness of the method. However, the observation that the most enriched sequences in the arsenic screenings are clearly distinct from those in the bismuth screenings suggests high sensitivity to the geometry of the linker atom. As the difference in the van der Waals radii between arsenic and bismuth is small,^43^ this effect might be primarily mediated by variations in metal-sulfur bond angles and lengths.

The four bicyclic peptides that were synthesised and investigated by SPR showed affinity for streptavidin in the low micromolar range, which aligns with previously reported peptides incorporating the HPQ motif.^32,44^ More importantly than the absolute affinities is the observation that the linear analogues exhibit two orders of magnitude lower affinity for streptavidin than their bicycles, despite the presence of the identical HPQ motif (**Figure 4**). This observation can be attributed to an entropic effect related to the preorganisation of the bicycle compared to the linear peptide and has been previously reported for peptide substrates of the Zika and West Nile virus proteases assessed in presence and absence of Bi(III).^17^

## Methods

### Design of phage-displayed library

The naïve unmodified M13 phage library used in this study was a gift from Prof. Ratmir Derda (University of Alberta). It encodes an ACX_4_CX_4_CGGGENLYFQS peptide at the N-terminus of recombinant pVIII on a type 88 phage vector that is similar to previously published work.^45^

### Modification of phage-displayed libraries

Amplified phage libraries were reduced using 2 mM TCEP-NaOH in sterile PBS (pH 7.4) for 30 min at room temperature. Using spin desalting columns, residual TCEP was removed *via* centrifugation at 2000 × *g* for 2.5 min. The reduced phage library solutions were then modified by exposure to either BiBr_3_ (50 mM in DMSO), gastrodenol or NaAsO_2_ (both 50 mM in sterile ultrapure water) at final concentrations of 120 μM for 5 min, before excess reagent was removed using spin desalting columns (2000 × *g*, 2.5 min).

### Biopanning of modified libraries

Biopanning campaigns were conducted against streptavidin-coated magnetic particles. Four selection rounds with increasing stringency were performed for each modified phage library (**Table S1**). For selection, a competitive elution protocol using 0.1 mM biotin was employed. Sanger and Nanopore sequencing were used to analyse the eluted phages. A portion of the remaining eluted phage was amplified in *Escherichia coli* ER2738 for subsequent selection rounds.

### In situ peptide bicyclisation

Purified linear peptide was solved in aqueous reducing buffer (20 mM HEPES-KOH pH 7.4, 150 mM NaCl, 2.5 mM TCEP) and exposed to 1.1 equivalents of gastrodenol or sodium arsenite at 1 mM, while the mixture was gently vortexed. Quantitative conversions of the linear peptides to the bicycles were confirmed with high-resolution mass spectrometry (**Figures S10 & S11**)

### Surface plasmon resonance

SPR measurements were performed with 20 mM HEPES-KOH pH 7.4, 150 mM NaCl, 0.5 mM TCEP, 0.05% (v/v) polysorbate 20 as running buffer. For each experiment, 100-150 μg streptavidin in 10 mM sodium acetate pH 3.9 buffer were freshly immobilised on a sensor chip using EDC/NHS chemistry. All binding experiments were conducted at 20 °C in single cycle kinetics mode, with the evaluation of bicyclic peptides conducted in duplicate (**Figure S14**).

## Conclusion

We present the first genetically encoded bicyclic peptide libraries containing bismuth or arsenic atoms in the peptide core. This method complements existing and well-established strategies using alkylating agents like TBMB. Bicycles form instantaneously with Bi(III), As(III) and potentially alternative thiophilic metal reagents in buffer at neutral pH. The use of water-soluble gastrodenol eliminates the need for previously necessary organic co-solvents. Modification of semi-randomised phage libraries proves highly reliable, as demonstrated by our similar results from repeated screenings. Bicyclic peptides exhibit dramatically increased affinity to the protein target compared to their linear analogues. We anticipate that this strategy will find broad applications, such as in the identification of bicyclic peptide ligands for radiopharmaceuticals.

## Acknowledgement

We thank the Australian Research Council for funding support, including a Discovery Project (DP230100079) and a Future Fellowship (FT220100010). We acknowledge Prof. Ratmir Derda (University of Alberta) for generously providing training and the unmodified phage library. We also thank Prof. Derda’s group members Dr. Alexey Atrazhev for sharing expertise in cloning and Dr. Arunika Ekanayake and Kejia Yan for valuable discussions. We also thank Dr. Claudia Yan at the Biomolecular Resource Facility (Australian National University) for performing Sanger sequencing, and Lavi Singh and Lydia Murphy in the group of A/Prof. Benjamin Schwessinger (Australian National University) for performing and supporting Nanopore sequencing. We further acknowledge Dr. Shouvik Aditya (Australian National University) for his assistance in SPR training and analysis as well as Richard Morewood (Australian National University) for training and assistance in HPLC purification.

