## Supporting Information for "Phage-Encoded Bismuth Bicycles: Instant Access to Targeted Bioactive Peptides"

**Abbreviations**

| | |
|---|---|
| DIPEA | *N*-ethyl-*N*-(propan-2-yl)propan-2-amine |
| DMF | dimethyl formamide |
| DMSO | dimethyl sulfoxide |
| DNA | deoxyribonucleic acid |
| dNTP | deoxynucleotide triphosphate |
| EDT | ethane-1,2-dithiol |
| EDTA | ethylenediaminetetraacetic acid |
| ESI | electron spray ionisation |
| FA | formic acid |
| Fmoc-AA | $N_\alpha$-fluorenylmethoxycarbonyl protected amino acid |
| HBTU | hexafluorophosphate benzotriazole tetramethyl uronium |
| HOBt | 1*H*-1,2,3-benzotriazol-1-ol |
| HPLC | high performance liquid chromatography |
| HRMS | high resolution mass spectrometry |
| HSA | human serum albumin |
| IPTG | isopropyl β-D-1-thiogalactopyranoside |
| LB | lysogeny broth |
| LC-MS | liquid chromatography mass spectrometry |
| PBS | phosphate-buffered saline |
| PCR | polymerase chain reaction |
| PEG | polyethylene glycol |
| pVIII | major coat protein |
| SPPS | solid-phase peptide synthesis |
| SPR | surface plasmon resonance |
| ssDNA | single strand deoxyribonucleic acid |
| TCEP | tris(2-carboxyethyl)phosphine |
| TFA | trifluoroacetic acid |
| TIPS | triisopropylsilane |
| X-Gal | 5-bromo-4-chloro-3-indolyl-β-D-galactopyranoside |

### Materials

The materials used in this study were obtained commercially from Sigma-Aldrich (USA), AK Scientific (USA), Thermo Fisher Scientific (USA) or New England Biolabs (USA), unless otherwise specified in the protocol. To prevent contamination phage or bacterial solutions were pipetted using sterile low-retention filter pipette tips from Neptune Scientific (Australia).

### Modification of phage-displayed library

Prior to the modification reaction, the amplified naïve 3-cysteine phage library (peptide insert $ACX_4CX_4CGGGENLYFQS$ on recombinant pVIII) was reduced using 2 mM TCEP. This reaction was performed by diluting 100 μL of the input phage library (see concentrations in biopanning section) with 880 μL of autoclaved PBS, which was followed by the addition of 20 μL of fresh 100 mM TCEP-NaOH (pH 7.5, in ultrapure water). This mixture was allowed to shake for 30 min at room temperature. The residual TCEP was removed using spin desalting columns (Thermo Fisher, 89883, USA) by centrifuging the resulting reduced phage library solution at 2000 × *g* for 2.5 min. The phage library solutions were then modified as outlined below. After each modification, excess reagent was removed using spin desalting columns as previously described.

*Modification with bismuth tribromide ($BiBr_3$)*

Bismuth tribromide ($BiBr_3$) in DMSO (50 mM) was added to the reduced phage solution while gently vortexed to a final volume of 1040 μL to obtain a final concentration of 120 μM bismuth(III). The reaction mixture was vortexed briefly once more and incubated at room temperature for 5 min.

*Modification with bismuth tripotassium dicitrate (gastrodenol)*

Bismuth tripotassium dicitrate (gastrodenol) in ultrapure sterile water (50 mM) was added to the reduced phage solution while gently vortexed to a final volume of 1040 μL to obtain a final concentration of 120 μM bismuth(III). The reaction mixture was vortexed briefly once more and incubated at room temperature for 5 min.

*Modification with sodium arsenite ($NaAsO_2$)*

Sodium arsenite ($NaAsO_2$) in ultrapure sterile water (50 mM) was added to the reduced phage solution while gently vortexed to a final volume of 1040 μL to obtain a final concentration of

120 µM arsenic(III). The reaction mixture was vortexed briefly once more and incubated at room temperature for 5 min.

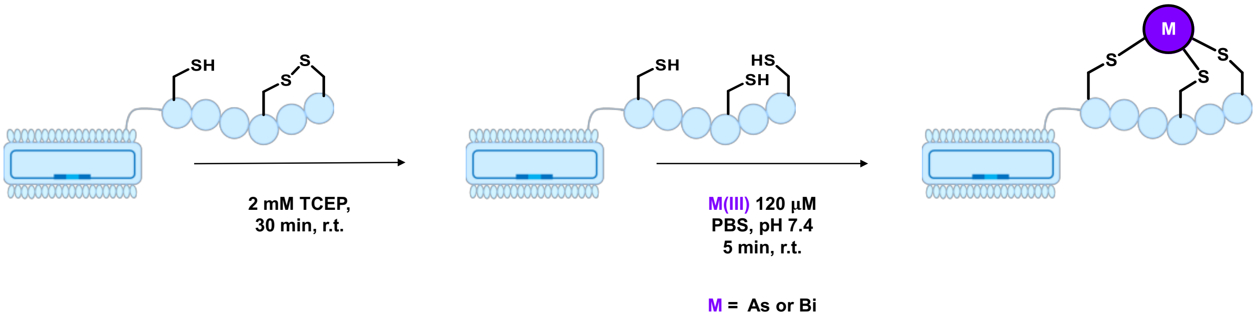

**Figure S1.** Phage library modification including reduction with TCEP and bicyclisation using arsenic(III) or bismuth(III).

### Biopanning campaigns of modified libraries

*Streptavidin biopanning campaign*

First, a 10 µL portion of streptavidin-coated paramagnetic particles (Promega, Z5481, USA) in a centrifuge tube was washed 3× with autoclaved PBS to remove the storage buffer using a magnetic stand, pulling away the beads in each washing step. Then, a blocking step was performed by incubating the washed beads in 100 µL of autoclaved PBS containing 0.5% (w/v) HSA for 2 h at room temperature. The blocking solution was then removed from the beads by magnetic separation followed by washing once with autoclaved PBS. Then, 100 µL of the modified (as described above) phage solution in PBS (~$10^{11}$ pfu/mL input phage concentration in round 1, ~ $10^{10}$ pfu/mL input phage concentration in round 2, ~$10^{9}$ pfu/mL input phage concentration in round 3 and ~$10^{8}$ pfu/ml input phage concentration in round 4 to increase the stringency) were added. The phage-bead mixture was incubated at room temperature with gentle shaking for 1 h. After the completion of 1 h incubation time, the beads were separated from the solution containing unbound phage particles and washed (3× in round 1, 6× in round 2, and 9× in rounds 3 and 4 to increase the stringency) with 100 µL of autoclaved PBS containing 0.05% (v/v) polysorbate 20 to ensure a more complete removal of unbound phage particles. Then, the competitive elution of the bound phage particles was carried out by suspending the beads in 100 µL of 0.1 mM biotin (in sterile water adjusted to pH 7.5 with 0.5 µL of 5 M NaOH solution) and incubating for 10 min at room temperature. The eluted phage particles were then removed from the beads by magnetic separation.

To determine the titre of recovered phage in each round, a 10 μL portion of the eluted phage solution was diluted 10- and 1000-fold in autoclaved PBS to perform a plaque-forming assay with these dilutions as described below.

**Table S1.** Increasing stringency in the selection rounds against immobilised streptavidin.

| Round | Magnetic beads [μL] | Phage input [pfu/mL] | Detergent [% v/v] | Washes |
|---|---|---|---|---|
| 1 | 10 | ~$10^{11}$ | 0.05 | 3 |
| 2 | 10 | ~$10^{10}$ | 0.05 | 6 |
| 3 | 10 | ~$10^{9}$ | 0.05 | 9 |
| 4 | 10 | ~$10^{8}$ | 0.05 | 9 |

**Plaque-forming assay**

An overnight culture of *E. coli* ER2738 (New England Biolabs, E4104, USA), a strain commonly used for phage display, was prepared by growing the bacteria in tetracycline containing LB media at 37 °C for 16 h at 220 rpm. Aliquots of 3 mL of melted top agar were transferred into sterile polystyrene tubes (Corning, 352051, USA) and heated up to 50 °C on a dry bath (Thermo Fisher, 88870004, USA) until ready to use. A dilution series of the phage library was prepared as required, at a final volume of 1 mL, whereby a 10 μL portion of each phage dilution was added to 200 μL of the bacterial overnight culture, incubated for 5 min and transferred fully into a tube containing 3 mL top agar. The contents in the tube were mixed gently by vortexing and poured entirely on a warm LB/IPTG/X-Gal plate, allowing the top agar-mixture to evenly spread out on the plate. The plate was allowed to solidify and left in an incubator oven overnight at 37 °C. The number of plaques was counted by selecting a plate with plaques separated at a countable level. The concentration of the original solution was then calculated in pfu/mL using the following equation.

Concentration of the original phage library solution = $\frac{number\ of\ plaques}{0.01}$ × dilution factor (pfu/mL)

**Amplification of phage eluents**

To amplify the phage eluents by infecting bacterial cells, a 70 µL portion of the eluted phage solution was used. This was carried out by transferring 250 µL of *E. coli* ER2738 overnight culture into a 250 mL baffled Erlenmeyer flask with 25 mL of LB media containing tetracycline, for 1 h at 37 °C at 220 rpm (to reach $OD_{600}$ values up to 0.4–0.5) followed by the addition of 70 µL of the phage eluent after the desired OD value was reached and allowing this bacterial/phage mixture to grow for 16 h at 30 °C at 200 rpm.

After growing the culture for 16 h, all the contents in the flask were transferred into a sterile 50 mL centrifuge tube (Corning, 352070, USA) and centrifuged at 5000 × *g* for 15 min at 4 °C. After the centrifugation, the supernatant was transferred into a new sterile 50 mL centrifuge tube containing 5 mL of cooled 6× PEG-NaCl and the solution was gently mixed by vortexing. The solution was then kept overnight at 4 °C to precipitate the phage particles.

The precipitated phage was then separated from the supernatant by centrifuging the 50 mL tubes at 14000 × *g* for 30 min at 4 °C. The supernatant was decanted into a bleach waste container. The tubes were kept upright for 15 min to help remove any residual liquid. Then the pellet was dissolved in 1 mL of autoclaved PBS and the solution was transferred into a 1.5 mL tube followed by centrifugation at 21000 × *g* at 4 °C for 2 min to remove any insoluble particles.

The supernatant was then transferred into a 1.5 mL tube containing 300 µL of cooled 6× PEG-NaCl. The solution was gently mixed by vortexing and incubated for 1 h on ice. After completion of the incubation, resulting in precipitation of the phage particles, the tubes were centrifuged at 14000 × *g* for 30 min at 4 °C to separate phage particles from the solution. The pellet was then dissolved in 1 mL of autoclaved PBS and the titer of the phage solution was determined by taking 10 µL of this amplified solution for preparing a dilution series of $10^2$, $10^4$, $10^6$, $10^{8,}$ and $10^{10}$ and performing the plaque-forming assay using $10^6$, $10^8$ and $10^{10}$ diluted solutions.

Finally, 200 µL of the amplified phage solution was kept for the biopanning of the next round by storing short-term at 4 °C while the remaining 800 µL phage solution was mixed with sterile glycerol solution (final glycerol content 50%) for medium-term storage at –20 °C.

**Preparation of Sanger sequencing samples**

For an initial analysis of the selection outcome, several plaques from the LB/IPTG/X-Gal plates from the plaque-forming assay were carefully picked to avoid cross contamination and added to a 50 mL centrifuge tube containing 5 mL LB containing tetracycline, and 50 μL of overnight *E. coli* ER 2738 culture and allowed to grow at 30 °C for 16 h at 200 rpm. The cells were then pelleted down by centrifuging at 5000 × g for 15 min at 4 °C followed by removal of the supernatant. The cells were lysed, and the phage ssDNA was extracted from the lysed bacterial cells by performing the mini prep protocol adapted from the manufacturer's protocol (New England Biolabs, T1010L, USA). To amplify the desired region of the phage plasmid, BigDye PCR was carried out and the obtained PCR products were purified for use in Sanger sequencing on an AB 3730xl DNA analyser (Thermo Fisher, USA) at the Biomolecular Resource Facility (John Curtin School of Medical Research, Australian National University).

*Extraction of phage ssDNA*

A commercially available miniprep set (New England Biolabs, T1010L, USA) was used for extraction and purification of phage ssDNA. The cell pellet was resuspended in 200 μL of cell resuspension buffer and vortexed to ensure the absence of visible clumps of cells. Then, 200 μL of lysis buffer was added to lyse the cells followed by incubation at room temperature for 1 min. 400 μL of the neutralisation buffer was then added to neutralise the lysate followed by incubating at room temperature for 1 min. Following the incubation, the samples were subjected to centrifugation at 16000 × *g* at room temperature for 10 min. The supernatant was then transferred into columns in 2 mL collection tubes followed by centrifugation at 16000 × *g* for 1 min at room temperature. The flow-through was discarded and 200 μL of wash buffer 1 were added to the column and centrifuged at 16000 × *g* for 1 min at room temperature. Then 400 μL of wash buffer 2 were added to the column and centrifuged at 16000 × *g* for 1 min at room temperature. The flow-through was discarded and the columns were centrifuged at 16000 × *g* for 1 min at room temperature to remove residual ethanol followed by placing the columns inside 1.5 mL tubes. The DNA was eluted by adding 30 μL of sterile ultrapure water at 55 °C, directly onto the silica membrane, incubating for 1 min at room temperature and centrifuging at 16000 × g for 1 min at room temperature.

*Sanger sequencing PCR*

PCR reactions were performed on a C1000 Touch Thermal Cycler (BioRad, USA).

Sanger PCR contents:

1. DNA template: 300–600 ng
2. sequencing buffer for BigDye (5×): 4 µL
3. BigDye: 1 µL
4. reverse primer (10 µM): 3.2 µL
5. nuclease-free water: fill to 20 µL

Thermocycler settings:

a. 94 °C, 5 min
b. 96 °C, 10 s
c. 50 °C, 5 s
d. 60 °C, 4 min
e. repeat b–d for 35 cycles
f. 4 °C, hold

*Sanger sequencing PCR product clean-up*

In a 1.5 mL tube, 5 µL of 125 mM EDTA were added to the 20 µL of the PCR product followed by the addition of 60 µL of 100% (v/v) EtOH. The contents of the tube were mixed by vortexing, spun down briefly, and incubated at room temperature for 15 min. Then, the tubes were centrifuged at 12000 × *g* for 20 min at room temperature. The supernatant was discarded carefully and 270 µL of 70% (v/v) EtOH in ultrapure water was added. The tubes were centrifuged again at 12000 × *g* for 10 min at room temperature. The resulting supernatant was discarded, and the tube dried under vacuum for 15 min and turned over for Sanger sequencing at the Biomolecular Resource Facility (John Curtin School of Medical Research, Australian National University). Sequencing results were analysed with Genious Prime 2023 (Dotmatics, USA) by aligning their reverse complement to the full phage genome and translating the peptide insert. The peptide inserts ($ACX_4CX_4CENLYFQS$) were deemed valid if they were 22 amino acids long and contained 3 or more cysteine residues.

**Sequencing primers**

Primers were obtained from Integrated DNA Technologies (Singapore).

**Table S2.** Primers used in this study for our M13 phage library on pVIII.

| Primer | Sequence | Length [bp] | $T_m$ [°C] |
|---|---|---|---|
| Forward | TATCCAAACAAAGAGTCATGAGC | 23 | 57.1 |
| Reverse | GAGTTTCGTCACCAGTACAAAC | 22 | 58.4 |

**Preparation of Nanopore sequencing samples**

Phage eluents or amplified phage solutions were amplified by performing PCR with Phusion polymerase (Thermo Fisher, USA). The obtained PCR products were purified by agarose gel purification following the respective manufacturer's protocol (Meridian Bioscience, 52060, USA). To increase the yield of obtained DNA, a two-step PCR protocol was implemented.

*Nanopore sequencing PCR*

Nanopore PCR step 1 contents:

1. DNA template (phage solution): 2 µL (amplified) or 15 µL (elution)
2. Phusion HF buffer (5×): 10 µL
3. Phusion polymerase (1 unit): 0.5 µL
4. dNTPs (10 mM): 1 µL
5. forward primer (10 µM): 2.5 µL
6. reverse primer (10 µM): 2.5 µL
7. nuclease-free water: fill to 50 µL

Thermocycler settings:

a. 98 °C, 3 min
b. 98 °C, 10 s
c. 50 °C, 20 s
d. 72 °C, 30 s
e. repeat b–d for 10 cycles
f. 98 °C, 10 s

g. 72 °C, 30 s
h. repeat f–g for 20 cycles
i. 72 °C, 5 min
j. 4 °C, hold

Nanopore PCR step 2 contents:

1. PCR product from step 1: 66–80 ng
2. Phusion HF buffer (5×): 10 μL
3. Phusion polymerase (1 unit): 0.5 μL
4. dNTPs (10 mM): 1 μL
5. forward primer (10 μM): 2.5 μL
6. reverse primer (10 μM): 2.5 μL
7. nuclease-free water: fill to 50 μL

Thermocycler settings:

a. 94 °C, 5 min
b. 96 °C, 10 s
c. 50 °C, 5 s
d. 60 °C, 4 min
e. repeat b–d for 35 cycles
f. 4 °C, hold

*Nanopore sequencing PCR product clean-up*

A commercially available PCR purification kit (Meridian Bioscience, 52060, USA) was used for purification of the PCR products for Nanopore sequencing. The PCR product samples were first mixed with 2 volumes of the provided binding buffer and loaded onto a column within a 2 mL collection tube followed by centrifugation at 11000 × *g* for 30 s at room temperature. The flow-through was discarded and 700 μL of the wash buffer were added to the column and centrifuged at 11000 × *g* for 30 s at room temperature. The flow-through was discarded and the step was repeated one more time to minimise chaotropic salt carry-over. The columns were then centrifuged at 11000 × *g* for 1 min at room temperature to remove residual ethanol. The columns were placed in 1.5 mL tubes and the DNA was eluted by adding 30 μL of autoclaved ultrapure water at 55 °C, directly onto the silica membrane, incubated for 1 min at room temperature and centrifuging at 11000 × *g* for 1 min at room temperature and turned over for Nanopore sequencing performed by the group of Prof. Benjamin Schwessinger (Research School of Biology, Australian National University) with the Nanopore Rapid Barcoding Kit 96 V14, on MinION R10.4.1 flowcells, using the MinION device from Oxford Nanopore Technologies, UK.

*Script for evaluation of Nanopore data*

A custom Python script for the extraction, ranking and visualisation of relevant displayed peptide sequences was written with assistance from Chat GPT-4 (OpenAI, USA) and will be made available upon request. Briefly, the script reads the raw *.fastq deep-sequencing file to find matching $ACX_4CX_4CGGGENLYFQS$ patterns in the amber readthrough translation of the phage DNA sequences, allowing up to three mismatches. The extracted sequences are then filtered to include only those with a length of 22 amino acids, no stop codons and at least 3 cysteine residues. These valid sequences are then ranked by their abundance and visualised in an image. Lastly, a dipeptide frequency analysis is performed with respect to the randomised $CX_4CX_4C$ part of the peptide insert. Excerpts from the logs regarding the sequencing data analysis are shown below.

**Table S3.** Summary of deep sequencing data analysis outcomes.

| Target | Library | Round | Total sequences | Valid pattern | Unique sequences |
|---|---|---|---|---|---|
| Streptavidin | $BiBr_3$ | 4 | 55764 | 39507 | 3049 |
| Streptavidin | gastrodenol | 4 | 85667 | 23153 | 5154 |
| Streptavidin | $NaAsO_2$ (1) | 4 | 78998 | 14133 | 1698 |
| Streptavidin | $NaAsO_2$ (2) | 4 | 65229 | 23009 | 3145 |

**Sanger sequencing results**

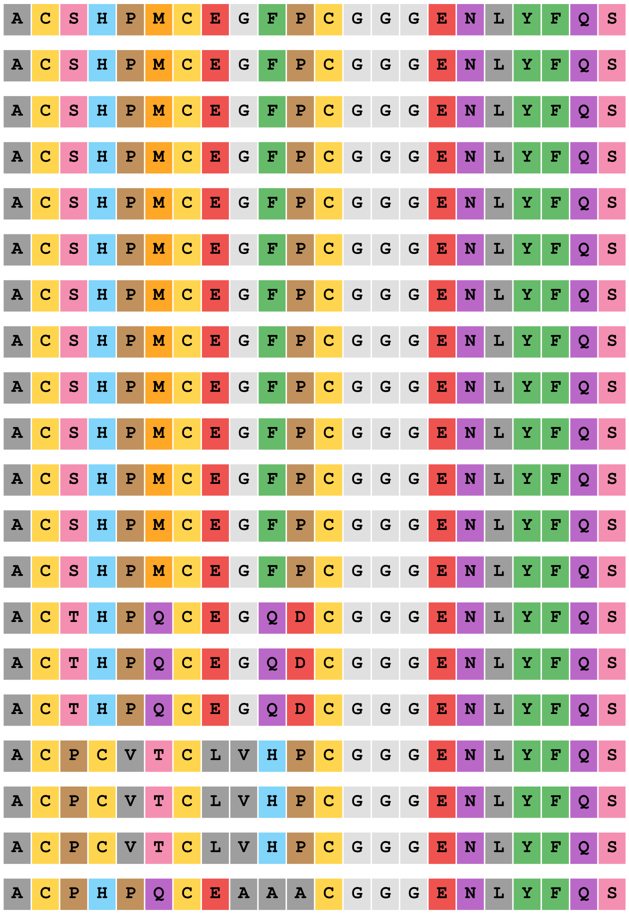

**Fig. S2.** Valid peptide inserts of 20 phage clones picked for Sanger sequencing from streptavidin $BiBr_3$ selection round 4.

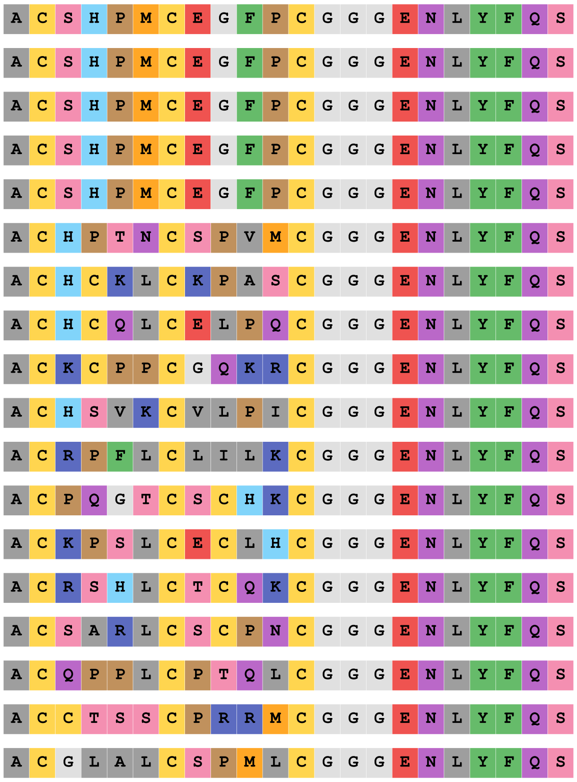

**Fig. S3.** Valid peptide inserts of 20 phage clones picked for Sanger sequencing from streptavidin gastrodenol selection round 4.

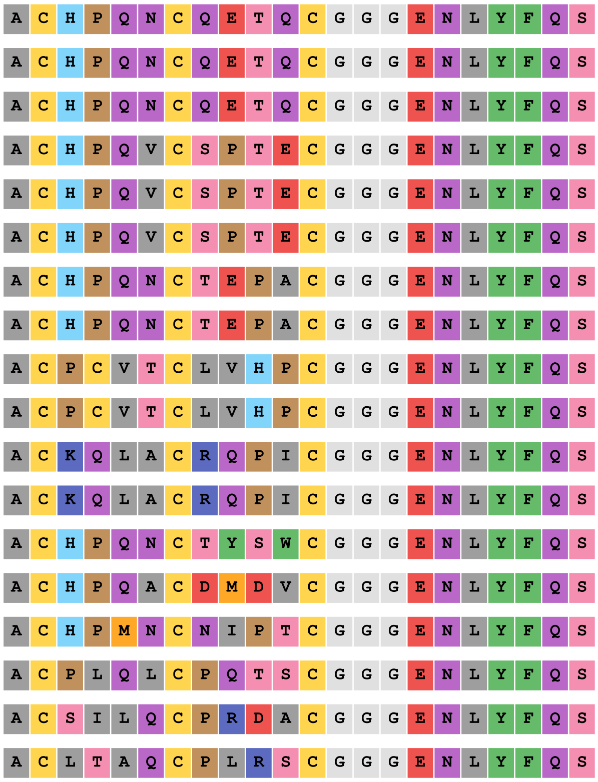

**Fig. S4.** Valid peptide inserts of 20 phage clones picked for Sanger sequencing from streptavidin $NaAsO_2$ selection (1) selection round 4.

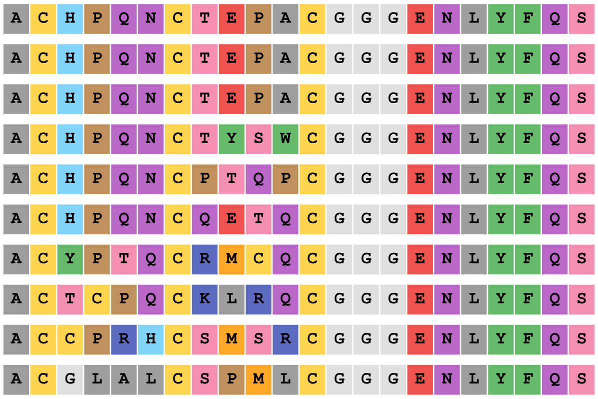

**Fig. S5.** Valid peptide inserts of 20 phage clones picked for Sanger sequencing from streptavidin $NaAsO_2$ selection (2) round 4.

### Nanopore sequencing results

| | | |
|---|---|---|
| #1 | A C S H P M C E G F P C G G G E N L Y F Q S | 37.73% |
| #2 | A C P C V T C L V H P C G G G E N L Y F Q S | 25.13% |
| #3 | A C T H P Q C E G Q D C G G G E N L Y F Q S | 6.44% |
| #4 | A C P H P Q C E A A A C G G G E N L Y F Q S | 5.67% |
| #5 | A C T T W G C T V Q H C G G G E N L Y F Q S | 0.90% |
| #6 | A C H P Q F C I S P E C G G G E N L Y F Q S | 0.81% |
| #7 | A C H V P P C R R K D C G G G E N L Y F Q S | 0.71% |
| #8 | A C Q Q L L C F R W I C G G G E N L Y F Q S | 0.58% |
| #9 | A C L G P T C N I Q N C G G G E N L Y F Q S | 0.35% |
| #10 | A C S H P M C E G F P C G G G E N L Y F Q L | 0.34% |
| #11 | A C H P Q N C T E P A C G G G E N L Y F Q S | 0.34% |
| #12 | A C P C V T C L V H P C G G G E N L Y F Q L | 0.31% |
| #13 | A C G L A L C S P M L C G G G E N L Y F Q S | 0.30% |
| #14 | A C S H P M C Q G F P C G G G E N L Y F Q S | 0.28% |
| #15 | A C R H P M C E G F P C G G G E N L Y F Q S | 0.27% |
| #16 | A C P L V K C P P C S C G G G E N L Y F Q S | 0.25% |
| #17 | A C H P Q F C S I T P C G G G E N L Y F Q S | 0.23% |
| #18 | A C H L H S C P E I Q C G G G E N L Y F Q S | 0.19% |
| #19 | A C S H P M C E G F P C G G G E N L Y S Q S | 0.16% |
| #20 | A C P S R Q C C N M Y C G G G E N L Y F Q S | 0.16% |
| #21 | A C H P Q V C S P T E C G G G E N L Y F Q S | 0.16% |
| #22 | A C H P Q F C Q T A P C G G G E N L Y F Q S | 0.15% |
| #23 | A C S H P M C E G F P C G G G E N L Y S S L | 0.15% |
| #24 | A C S H P M C E G F P C G G G E N L Y F H L | 0.13% |
| #25 | A C S H P M C E G F P C G G G E N L Y F P S | 0.13% |

**Fig. S6.** Top 25 peptide inserts ranked by percent abundance from Nanopore sequencing data of phages from streptavidin $BiBr_3$ selection round 4 as analysed by the Python script.

| | | | | | | | | | | | | | | | | | | | | | | |
|---|---|---|---|---|---|---|---|---|---|---|---|---|---|---|---|---|---|---|---|---|---|---|
| #1 | A | C | S | H | P | M | C | E | G | F | P | C | G | G | G | E | N | L | Y | F | Q | S | 18.64% |
| #2 | A | C | G | L | A | L | C | S | P | M | L | C | G | G | G | E | N | L | Y | F | Q | S | 3.79% |
| #3 | A | C | P | H | P | Q | C | E | A | A | A | C | G | G | G | E | N | L | Y | F | Q | S | 2.95% |
| #4 | A | C | H | C | K | L | C | K | P | A | S | C | G | G | G | E | N | L | Y | F | Q | S | 2.14% |
| #5 | A | C | Q | Q | L | L | C | F | R | W | I | C | G | G | G | E | N | L | Y | F | Q | S | 2.09% |
| #6 | A | C | T | H | P | Q | C | E | G | Q | D | C | G | G | G | E | N | L | Y | F | Q | S | 2.02% |
| #7 | A | C | K | T | Q | P | C | K | C | S | P | C | G | G | G | E | N | L | Y | F | Q | S | 1.90% |
| #8 | A | C | C | T | S | S | C | P | R | R | M | C | G | G | G | E | N | L | Y | F | Q | S | 1.12% |
| #9 | A | C | P | C | V | T | C | L | V | H | P | C | G | G | G | E | N | L | Y | F | Q | S | 0.98% |
| #10 | A | C | E | Y | E | T | C | R | Q | S | Q | C | G | G | G | E | N | L | Y | F | Q | S | 0.88% |
| #11 | A | C | S | A | Q | S | C | L | G | E | N | C | G | G | G | E | N | L | Y | F | Q | S | 0.56% |
| #12 | A | C | K | T | R | S | C | H | R | R | L | C | G | G | G | E | N | L | Y | F | Q | S | 0.54% |
| #13 | A | C | K | C | P | P | C | G | Q | K | R | C | G | G | G | E | N | L | Y | F | Q | S | 0.52% |
| #14 | A | C | A | N | S | W | C | C | K | A | P | C | G | G | G | E | N | L | Y | F | Q | S | 0.51% |
| #15 | A | C | L | T | A | Q | C | P | L | R | S | C | G | G | G | E | N | L | Y | F | Q | S | 0.49% |
| #16 | A | C | H | P | P | W | C | D | Y | R | F | C | G | G | G | E | N | L | Y | F | Q | S | 0.48% |
| #17 | A | C | P | S | R | Q | C | C | N | M | Y | C | G | G | G | E | N | L | Y | F | Q | S | 0.45% |
| #18 | G | S | R | Q | L | Q | C | C | P | A | K | C | G | G | G | E | N | L | Y | F | Q | S | 0.43% |
| #19 | A | C | H | K | R | S | C | C | A | S | A | C | G | G | G | E | N | L | Y | F | Q | S | 0.42% |
| #20 | A | C | P | L | V | K | C | P | P | C | S | C | G | G | G | E | N | L | Y | F | Q | S | 0.42% |
| #21 | A | C | Y | P | P | P | C | L | Q | N | L | C | G | G | G | E | N | L | Y | F | Q | S | 0.38% |
| #22 | A | C | R | T | S | Q | C | S | C | N | H | C | G | G | G | E | N | L | Y | F | Q | S | 0.33% |
| #23 | A | C | S | L | Q | C | C | P | F | R | Q | C | G | G | G | E | N | L | Y | F | Q | S | 0.31% |
| #24 | A | C | H | P | Q | N | C | T | E | P | A | C | G | G | G | E | N | L | Y | F | Q | S | 0.31% |
| #25 | A | C | G | P | T | P | C | E | I | Q | L | C | G | G | G | E | N | L | Y | F | Q | S | 0.29% |

**Fig. S7.** Top 25 peptide inserts ranked by percent abundance from Nanopore sequencing data of phages from streptavidin gastrodenol selection round 4 as analysed by the Python script.

| | | |
|---|---|---|
| #1 | A C H P Q N C T E P A C G G G E N L Y F Q S | 14.24% |
| #2 | A C P C V T C L V H P C G G G E N L Y F Q S | 13.03% |
| #3 | A C H P Q N C Q E T Q C G G G E N L Y F Q S | 12.44% |
| #4 | G S R Q L Q C C P A K C G G G E N L Y F Q S | 8.04% |
| #5 | A C K Q L A C R Q P I C G G G E N L Y F Q S | 3.98% |
| #6 | A C H P Q V C S P T E C G G G E N L Y F Q S | 3.96% |
| #7 | A C R C E H C P W P Q C G G G E N L Y F Q S | 2.85% |
| #8 | A C H P Q V C D R Q M C G G G E N L Y F Q S | 2.00% |
| #9 | A C H P Q N C T Y S W C G G G E N L Y F Q S | 1.95% |
| #10 | A C H P Q A C S Y S E C G G G E N L Y F Q S | 1.89% |
| #11 | A C H P Q N C T M N Q C G G G E N L Y F Q S | 1.78% |
| #12 | A C E T E V C S V G N C G G G E N L Y F Q S | 1.75% |
| #13 | A C T G P A C L T S T C G G G E N L Y F Q S | 1.28% |
| #14 | A C H P Q N C P T Q P C G G G E N L Y F Q S | 1.15% |
| #15 | A C H P Q N C A G T V C G G G E N L Y F Q S | 1.03% |
| #16 | A C H P Q V C H P V A C G G G E N L Y F Q S | 0.71% |
| #17 | A C G L A L C S P M L C G G G E N L Y F Q S | 0.63% |
| #18 | A C P L Q L C P Q T S C G G G E N L Y F Q S | 0.55% |
| #19 | A C H P M N C N I P T C G G G E N L Y F Q S | 0.53% |
| #20 | A C P I L K C I Q L Q C G G G E N L Y F Q S | 0.52% |
| #21 | A C H P Q V C S P Q R C G G G E N L Y F Q S | 0.49% |
| #22 | A C H P Q V C S T F A C G G G E N L Y F Q S | 0.46% |
| #23 | A C H P Q V C S H T L C G G G E N L Y F Q S | 0.46% |
| #24 | A C S H P M C E G F P C G G G E N L Y F Q S | 0.44% |
| #25 | A C H P Q N C P Q E P C G G G E N L Y F Q S | 0.36% |

**Fig. S8.** Top 25 peptide inserts ranked by percent abundance from Nanopore sequencing data of phages from streptavidin $NaAsO_2$ selection (1) round 4 as analysed by the Python script.

| | | |
|---|---|---|
| #1 | A C H P Q N C T E P A C G G G E N L Y F Q S | 29.63% |
| #2 | A C G L A L C S P M L C G G G E N L Y F Q S | 6.70% |
| #3 | A C H P Q V C S P T E C G G G E N L Y F Q S | 3.67% |
| #4 | A C H P Q N C Q E T Q C G G G E N L Y F Q S | 3.63% |
| #5 | A C H P Q N C T Y S W C G G G E N L Y F Q S | 2.81% |
| #6 | A C H P Q N C P T Q P C G G G E N L Y F Q S | 2.05% |
| #7 | A C H P Q V C D R Q M C G G G E N L Y F Q S | 1.49% |
| #8 | A C H P Q N C T M N Q C G G G E N L Y F Q S | 1.46% |
| #9 | A C R K D Y C Q Q P F C G G G E N L Y F Q S | 1.16% |
| #10 | A C Q R P C C Y Q Q P C G G G E N L Y F Q S | 1.00% |
| #11 | G S R Q L Q C C P A K C G G G E N L Y F Q S | 0.96% |
| #12 | A C P C V T C L V H P C G G G E N L Y F Q S | 0.82% |
| #13 | A C Q Q L L C F R W I C G G G E N L Y F Q S | 0.76% |
| #14 | A C T Q P A C G Q D Y C G G G E N L Y F Q S | 0.71% |
| #15 | A C H P Q N C L A P S C G G G E N L Y F Q S | 0.62% |
| #16 | C G G S R P C P R L A C G G G E N L Y F Q S | 0.60% |
| #17 | A C P P P L C L P R T C G G G E N L Y F Q S | 0.60% |
| #18 | A C H P Q N C A G T V C G G G E N L Y F Q S | 0.59% |
| #19 | A C P S R Q C C N M Y C G G G E N L Y F Q S | 0.56% |
| #20 | A C Q P W T C T R G K C G G G E N L Y F Q S | 0.51% |
| #21 | A C H P Q A C S Y S E C G G G E N L Y F Q S | 0.49% |
| #22 | A C S D S L C Q T V Q C G G G E N L Y F Q S | 0.47% |
| #23 | A C V N P T C E R Y K C G G G E N L Y F Q S | 0.42% |
| #24 | A C S Q S H C P S A Y C G G G E N L Y F Q S | 0.41% |
| #25 | A C P T V A C R T S Q C G G G E N L Y F Q S | 0.40% |

**Fig. S9.** Top 25 peptide inserts ranked by percent abundance from Nanopore sequencing data of phages from streptavidin $NaAsO_2$ selection (2) round 4 as analysed by the Python script.

### General conditions for linear peptide synthesis

Automated SPPS was performed using the Initiator+ Alstra peptide synthesiser (Biotage, Sweden). High-loading (GL Biochem, China or Auspep, Australia) or low-loading (Novabiochem, USA) Rink amide resins were used for SPPS. Reagents were used from commercial sources without further purification: Fmoc-AA (GL Biochem, China), coupling reagents (GL Biochem, China), solvents (ChemSupply, Australia), remaining reagents (Sigma Aldrich, USA or AK Sci, USA).

| | |
|---|---|
| *Resin swelling* | $CH_2Cl_2$, 60 min. |
| *Standard washing* | DMF, 5 min. |
| *Fmoc deprotection* | Piperidine in DMF (20% v/v), 1× 3 min, 1× 10 min. |
| *Amino acid coupling* | HBTU (3 eq), HOBt (3 eq), DIPEA (4 eq), Fmoc-AA (3 eq) in DMF, room temperature, 1× or 2× 1 h. |
| *Final washing* | $CH_2Cl_2$, 15 min. |
| *Resin cleavage* | TFA, thioanisole, EDT, TIPS, water (85:5:5:3:2 v/v), 2 h. |
| *Work-up* | Precipitation of entire cleavage solution in ice-cold $Et_2O$. Centrifugation at 4000 × *g*, 10 min. High-vacuum drying for 1 h. |

### Linear peptide sequences

**Table S4.** Linear peptide sequences synthesised in this study.

| Compound | N-terminus | Sequence | C-terminus |
|---|---|---|---|
| **1** | H- | A<u>C</u>SHPM<u>C</u>EGFP<u>C</u> | $-NH_2$ |
| **2** | H- | A<u>C</u>PHPQ<u>C</u>EAAA<u>C</u> | $-NH_2$ |
| **3** | H- | A<u>C</u>HPQN<u>C</u>TEPA<u>C</u> | $-NH_2$ |
| **4** | H- | A<u>C</u>HPQV<u>C</u>SPTE<u>C</u> | $-NH_2$ |

**Linear peptide purification**

Linear peptides underwent purification on an HPLC 600 system (Waters, USA) using preparative reverse phase HPLC. Isolated fractions were lyophilised using an Alpha 1-2 freeze dryer (Christ, Germany).

*Preparative HPLC methods*

Method A: This method was employed for the purification of peptides **1**, **3** and **4** utilising column 3. The gradient elution system utilised in this procedure comprises two solvents, ultrapure water and methanol, both containing 0.1% TFA as additive. Operating at a flow rate of 10.0 mL/min, the gradient commenced with 20% B in A and gradually increased to 90% B in A between minute 5 and 20. The proportion of B remained constant until the end of the purification process at minute 40.

Method B: This method was employed for the purification of **2** utilising column 4. The gradient elution system utilised in this procedure comprises two solvents, ultrapure water and methanol, both containing 0.1% TFA as additive. Operating at a flow rate of 10.0 mL/min, the concentration of solvent B commenced at 0% and progressively increased to 50% between minute 5 and minute 20. The proportion of solvent B remained constant until the purification process ended at minute 40.

*Columns*

Column 1: Eclipse XDB-$C_{18}$, 2.1 mm × 50 mm, 1.8 µm

Column 2: Alltima HP, $C_{18}$-AQ, 2.1 mm × 150 mm, 5 µm

Column 3: SymmetryPrep $C_{18}$, 19 mm × 150 mm, 7 µm

Column 4: YMC-Pack ODS-A $C_{18}$, 50 mm × 250 mm, 5 µm

**HRMS analysis**

Peptides were analysed using high-resolution ESI+ mass spectrometry with a Synapt G2-Si mass spectrometer (Waters, USA). Expected ion masses and isotope patterns were calculated with enviPat (Eawag, Switzerland).

**Table S5.** High-resolution mass spectrometry data of linear and bicyclic peptides.

| Compound | Molecular formula | Ion | Calculated | Observed |
|---|---|---|---|---|
| **1** | $C_{52}H_{77}N_{15}O_{15}S_4$ | $[M+K]^+$ | 1318.4238 | 1318.4248 |
| **1b** | $C_{52}H_{74}BiN_{15}O_{15}S_4$ | $[M+K]^+$ | 1524.3807 | 1524.3779 |
| **2** | $C_{47}H_{74}N_{16}O_{15}S_3$ | $[M+Na]^+$ | 1221.4574 | 1221.4568 |
| **2b** | $C_{47}H_{71}BiN_{16}O_{15}S_3$ | $[M+K]^+$ | 1443.3882 | 1443.3912 |
| **3** | $C_{49}H_{77}N_{17}O_{17}S_3$ | $[M+Na]^+$ | 1294.4738 | 1294.4747 |
| **3a** | $C_{49}H_{74}AsN_{17}O_{17}S_3$ | $[M+H]^+$ | 1344.3899 | 1344.3901 |
| **4** | $C_{50}H_{80}N_{16}O_{17}S_3$ | $[M+H]^+$ | 1273.5122 | 1273.5127 |
| **4a** | $C_{50}H_{77}AsN_{16}O_{17}S_3$ | $[M+K]^+$ | 1383.3662 | 1383.3673 |

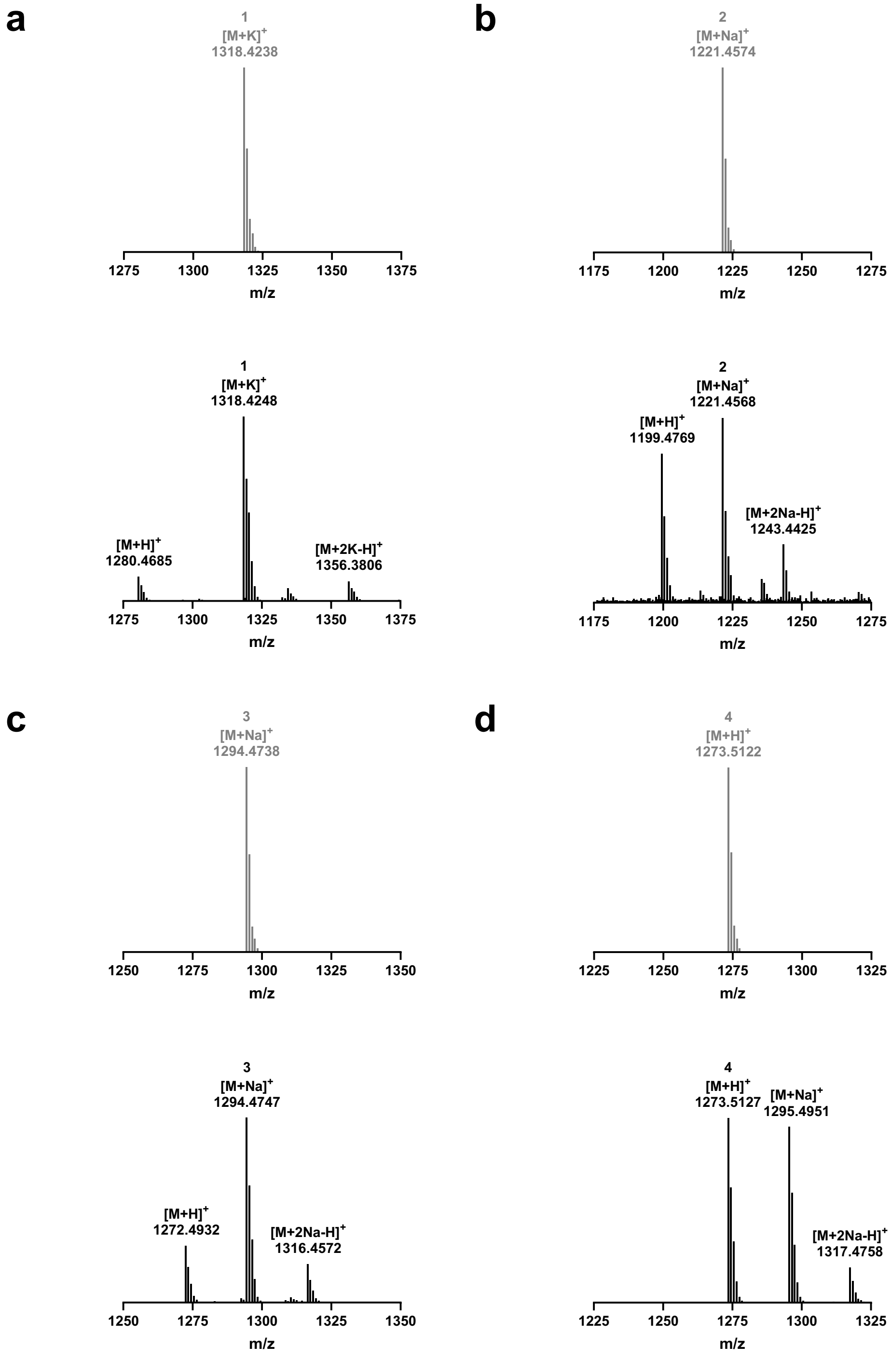

**Fig. S10.** Simulated isotope patterns (grey) and high-resolution mass spectra (black) of linear peptides (a) **1**, (b) **2**, (c) **3**, (d) **4**.

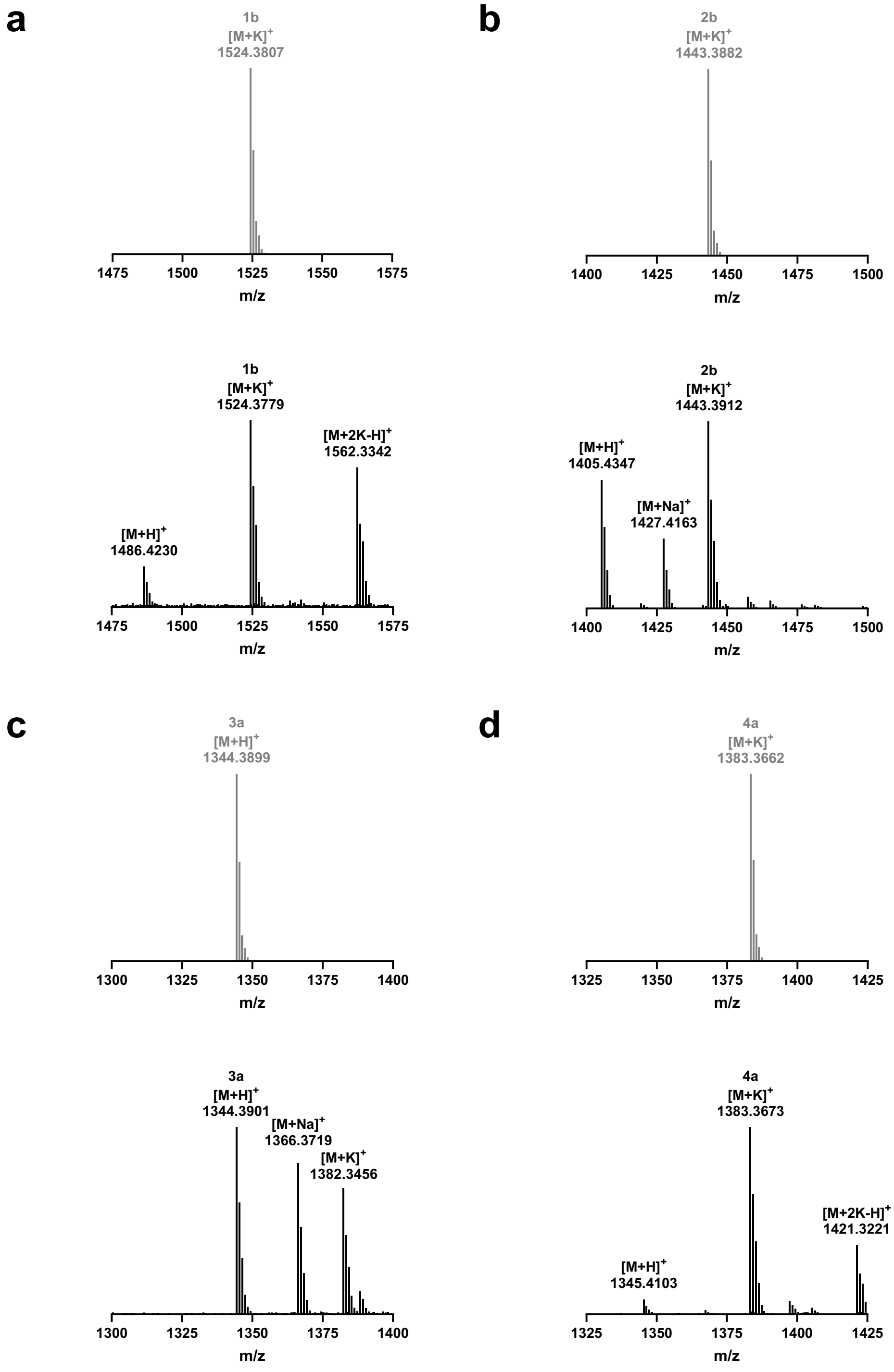

**Fig. S11.** Simulated isotope patterns (grey) and high-resolution mass spectra (black) of bicyclic peptides (a) **1b**, (b) **2b**, (c) **3a**, (d) **4a**.

**LC-MS analysis**

Identity of the linear and bismuth-peptide bicycles was confirmed using the analytical reverse phase LC-MS system 1260/6120 (Agilent, USA).

*Analytical LC-MS methods*

Method C: This gradient system utilised two solvents, A and B, namely ultrapure water (additive: 0.1% FA) and acetonitrile (additive: 0.1% TFA). Operating at a flow rate of 0.3 mL/min and using column 1, the gradient commenced with 20% B in A for the initial 2 min and gradually escalated to 95% over the subsequent 20 min. Subsequently, the proportion of B rapidly dropped to 10% within 1 min and this composition was maintained for 5 min.

Method D: This gradient system utilised two solvents, A and B, namely ultrapure water (additive: 0.1% FA) and acetonitrile (additive: 0.1% TFA). Operating at a flow rate of 1.0 mL/min and using column 2, the gradient commenced with 100% A for the initial 0.01 min and gradually escalated to 50% B over the subsequent 20 min. Subsequently, the proportion of B in A rapidly dropped to 0% within 1 min and this composition was maintained for 5 min.

*LC-MS traces and spectra*

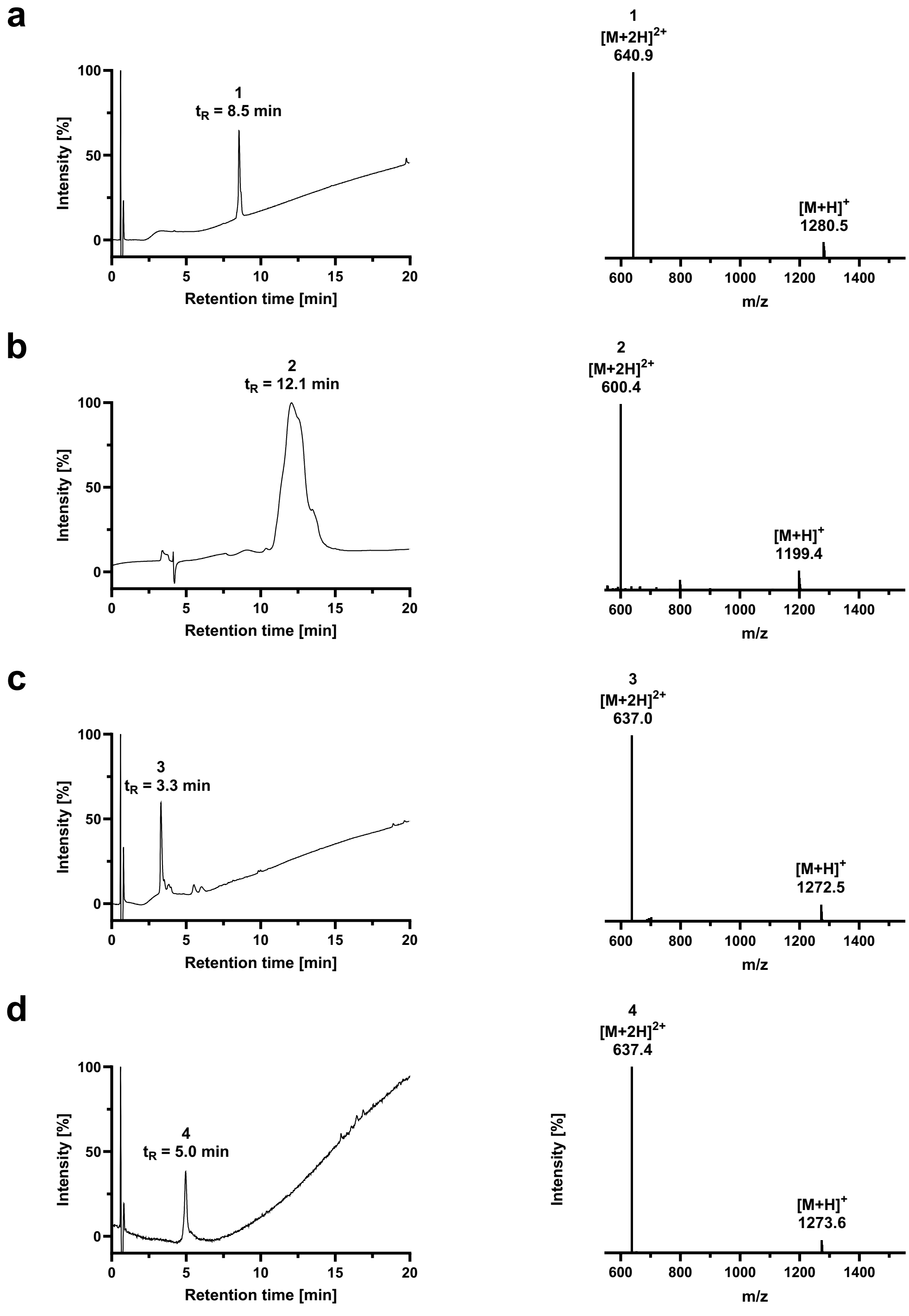

**Fig. S12.** LC-MS traces (214 nm) and spectra (ESI) of linear peptides (a) **1** (method C), (b) **2** (method D), (c) **3** (method C), (d) **4** (method C).

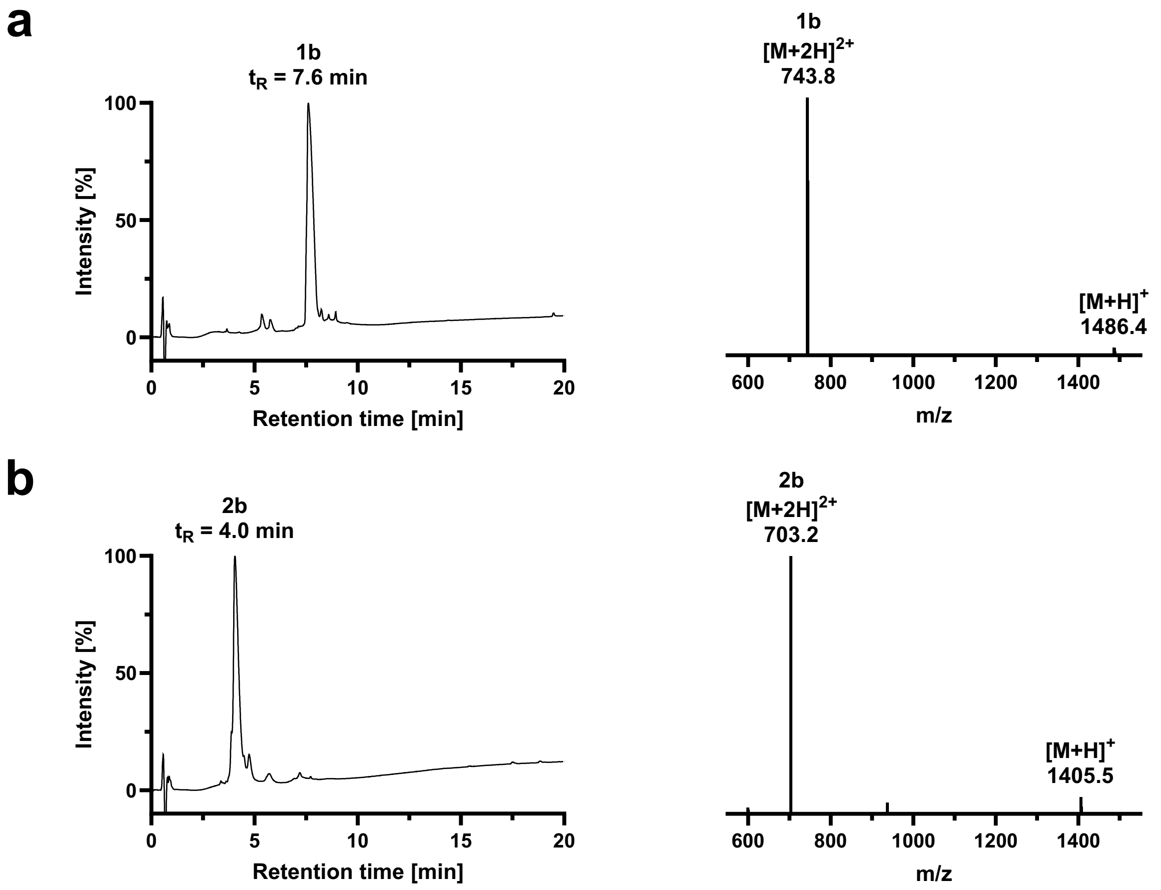

**Fig. S13.** LC-MS traces (214 nm) and spectra (ESI) of bismuth-peptide bicycles (a) **1b** (method C) and (b) **2b** (method C) prepared *in situ* with gastrodenol.

### SPR analysis

SPR experiments were performed on a Biacore 8K (Cytiva, USA) using a CM5 chip (Cytiva, 29104988, USA), onto which 100-150 µg streptavidin (New England Biolabs, USA) were freshly immobilised using an amine coupling kit (Cytiva, USA). The data were plotted and analysed in GraphPad Prism 10 (Dotmatics, USA) using the non-linear fit 'one site – specific binding'.

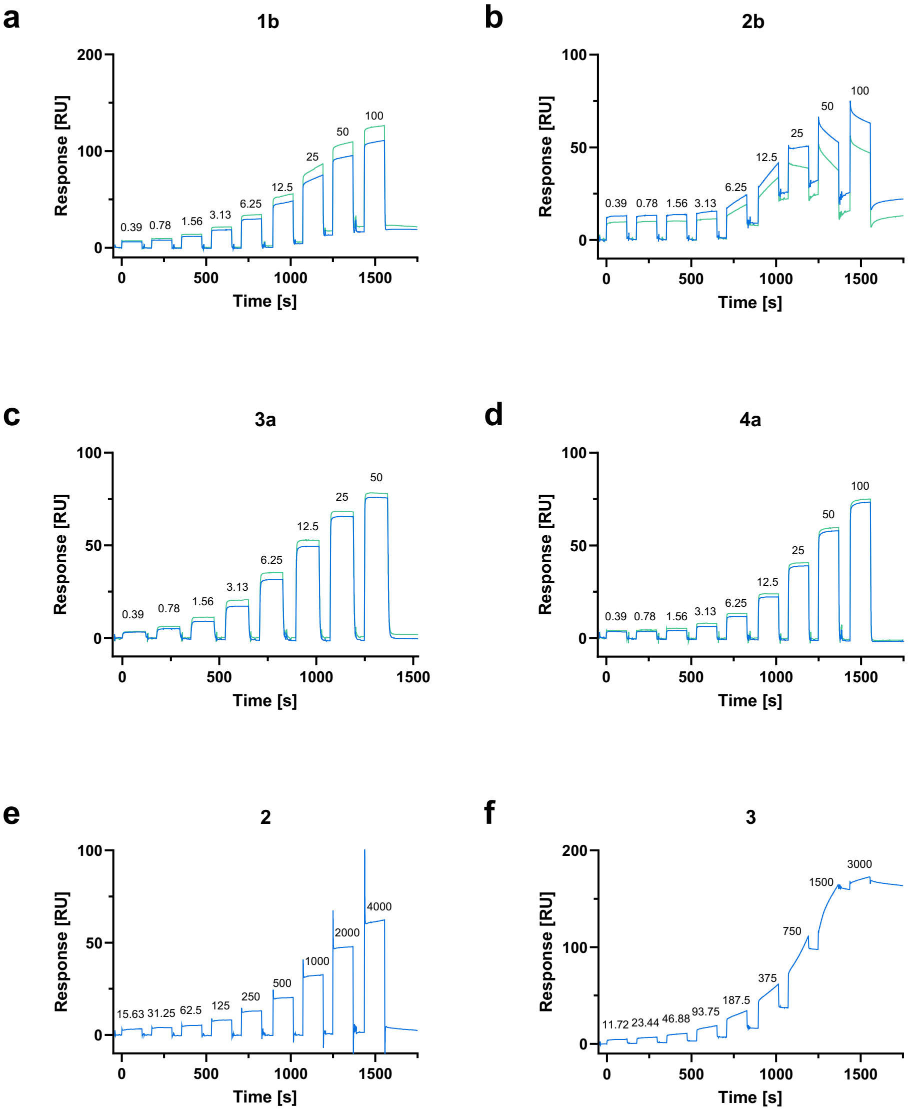

**Fig. S14.** SPR sensograms showing the binding response of peptides (a) **1b**, (b) **2b**, (c) **3a**, (d) **4a**, (e) **2** and (f) **3** to streptavidin immobilised on a CM5 chip in single kinetics mode. Rounded concentrations (in µM) are indicated above the response peaks. Individual replicates for bicyclic peptides are indicated in different colours (blue and green).

**Preparation of selected reagents for phage display**

*LB liquid medium*

10 g tryptone

5 g yeast extract

5 g sodium chloride

Fill to 1 L with deionised water.

Autoclave and store at room temperature.

*X-Gal/IPTG stock*

0.5 g IPTG

0.4 g X-Gal

Fill to 10 mL with DMF.

Store at –20 °C.

*LB/X-Gal/IPTG agar plates*

10 g tryptone

5 g yeast extract

5 g sodium chloride

15 g agar

Fill to 1 L with deionised water.

Autoclave and cool to 50 °C, then add 1 mL X-Gal/IPTG stock in DMF.

Pour into petri dishes, store at 4 °C.

*Top agar*

2 g tryptone

1 g yeast extract

1 g sodium chloride

1.5 g agar

Fill to 200 mL with deionised water.

Autoclave, store at room temperature and melt using microwave before use.

*6× PEG-NaCl*

90 g polyethylene glycol 8000 powder

52.6 g NaCl

Fill to 300 ml with autoclaved ultrapure water.

Autoclave in >1 L container to avoid spillage, allow for layers to mix, store at room temperature.
